# Aversive View Memory and Navigational Risk Sensitivity in the Desert Ant, *Cataglyphis Velox*

**DOI:** 10.1101/2021.09.24.461543

**Authors:** Cody A Freas, Antoine Wystrach, Sebastian Schwarz, Marcia L Spetch

## Abstract

Many ants establish foraging routes through learning views of the visual panorama. Route models have focused primarily on attractive view use, which experienced foragers orient towards to return to known sites. However, aversive views have recently been uncovered as a key component of route learning. Here, *Cataglyphis velox* rapidly learned aversive views, when associated with a negative outcome, a period of captivity in brush, triggering an increase in hesitation behavior. These memories were based on the accumulation of experiences over multiple trips with each new experience regulating forager’s hesitancy. Foragers were also sensitive to captivity time differences, suggesting they possess some mechanism to quantify duration. Finally, we analyzed foragers’ perception of risky (i.e. variable) versus stable aversive outcomes by associating two sites along the route with distinct captivity schedules, a fixed or variable duration, with the same mean across training. Foragers exhibited significantly less hesitation to the risky outcome compared to the fixed, indicating they perceived risky outcomes as less severe. Results align with a logarithmic relationship between captivity duration and hesitations, suggesting that aversive stimulus perception is a logarithm of its actual value. We conclude by characterizing how these behaviors can be executed within the mushroom bodies’ neural circuitry.

## Introduction

The navigational abilities of solitarily foraging ants can be attributed to a toolkit comprised of multiple strategies (Wehner et al. 1996; Collett et al. 2013; Cheng et al. 2014; Freas and Schultheiss 2018). The most well studied components of this toolkit are the path integration (PI) system and learned visual cues of the panorama. The PI system is informed via the celestial compass and a step-counter, creating a vector that continuously updates an estimate of the nest location during the foraging trip (Wehner 2003, 2008). PI can be especially useful for foragers when their environment lacks prominent terrestrial cues (Wehner 2020) or during a naïve forager’s first few foraging trips as they learn the surrounding terrestrial cues and form foraging routes (Kohler and Wehner 2005; Muller and Wehner 2010; Mangan and Webb 2012; Schwarz et al. 2017).

### View learning

When available, many ant species rapidly learn visual landmark information to navigate. Rather than attend to individual landmarks, foragers learn panoramic views around goal locations and along their foraging routes (Graham and Cheng 2009; Wystrach et al. 2011a). View learning first occurs around the nest before the onset of foraging, during learning walks (Wehner et al. 2004; Zeil and Fleischmann 2019) Foragers also acquire views en-route as they move away from known locations (Graham and Cheng 2009; Wystrach et al. 2011a; Schultheiss et al. 2016; Freas et al. 2018), and can rapidly learn the panorama at a new site, often after only one previous experience (Freas and Cheng 2018; Freas and Spetch 2019). Foragers retain long term-memories of these panoramas (Narendra et al. 2007) and, while navigating, compare these memories to their current view to recover their goal direction (Zeil 2012).

### Aversive views

View memories and their importance in route following have been well modeled (Zeil et al. 2003; Wystrach et al. 2011b; Baddeley et al. 2012; Kodzhabashev and Mangan 2015; Möller 2012), yet these models rely principally on the forager orienting towards attractive views via view comparison. Recent work has expanded this modelling to include the use of learned views that are repellant and cause foragers to turn away from views not associated with the current goal, resulting in orientation away from incorrect directions (Le Möel and Wystrach 2020; Murray et al. 2020). The interaction between these learned attractive and repellant views permits navigators to compare a single current view to their view memories to quickly decide whether to move toward or turn away from a given direction (Le Möel and Wystrach 2020, Murray et al. 2020). It has been suggested that learned views can become repellant depending on their orientation relative to the nest (Jayatikala et al. 2017; Murray et al. 2020), on the foraging motivational context (Schwarz et al. 2020) or when these views are associated with aversive outcomes (Wystrach et al. 2020). Previously attractive views can also become aversive when they are associated with negative outcomes. In both *Cataglyphis* and *Melophorus* desert ants, when a pit trap was added along a forager’s homeward route, resulting in foragers falling into brush, ants quickly memorized the views experienced just before this negative experience as repulsive. Eventually, after a few experiences falling into this pit, foragers formed new routes detouring around it. The interplay of aversive and attractive views appears to facilitate the formation of these detours. As foragers attempt to avoid aversive views, novel views that pilot around these areas become positively reinforced, leading to the development of new routes detouring around obstacles and areas with difficult terrain (Wystrach et al. 2020). Vegetation can often be hard for desert ants to move through effectively, especially when carrying food, resulting in increases to both expenditure of effort and the delay to return to the nest. After only a few trips experiencing the pit, foragers began to hesitate near the pit’s edge, increasing their hesitancy to pass through the area, evidenced through increases in both scanning behavior and path meander (Wystrach et al. 2020). Scanning behavior consists of a forager stopping forward movement and rotating their body on the spot. This behavior is associated with instances of increased navigational uncertainty: when the familiarity of the panorama decreases, the PI and panorama enter into conflict, or when the current route’s panorama is associated with failure (Wystrach et al. 2014, 2019). Thus, the incidence of scanning is a good behavioral proxy to assess the ant’s uncertainty and in this study we used it to quantify the strength of the aversion associated with a given location (Schwarz. et al. 2020).

### Risk perception

While navigating, foraging animals must make decisions assessing risky or safe options both in the resources they collect and in their foraging routes. Based on predictions within evolutionary theory, individuals were conventionally thought to make foraging choices that were strictly optimal, maximizing their net energy gains to increase fitness (MacArthur and Pianka 1966; Krebs 1986). However, humans and other animals sometimes behave in seemingly non-optimal or irrational ways with regards to their perception and preference for risk (Kahneman and Tversky 1979; Kacelnik and Bateson 1996). For example, when risk is generated by variability in amount, animals are often risk-averse or risk neutral, whereas when risk is generated by variability in delay, animals are typically risk-prone (e.g., Kacelnik and Bateson 1996; Kacelnik and Abreu 1998). Such irrational preferences are believed to flow from animals’ perception of the world, where true stimulus strength has a logarithmic relationship with the animal’s perception (Fechner 1860; Bruce and Johnson 1996; Kacelnik and Abreu 1998; Stevens and Marks 2017). Based on this principle, animals’ choices between risky (variable) and fixed outcomes should be predicted not by the mean value of these options but instead by their geometric means.

Risk perception and preference have been studied across a range of animals (for review see: Kacelnik and Bateson 1996; Kacelnik and El Mouden 2013). Much of the risk preference research in Hymenoptera has focused on foragers’ preference for risk solely in regards to the amount or quality of a given reward (Kacelnik and Bateson 1996; Hübner and Czaczkes 2017; De Agrò et al. 2021). In honeybees and bumble bees, a variety of outcomes have been reported with foragers showing evidence of no preference, risk avoidance and risk seeking foraging choices based on factors such as colony resource levels (Waddington et al. 1981; Cartar 1991; Perez and Waddington 1996; Fülöp and Menzel 2000). In ants, risk perception and sensitivity have been explored on the colony level, focusing on how collective decision-making influences choice in the assessment of potential nesting site quality and in food reward quality. Rock ants (*Temnothorax albipennis*) were shown to exhibit risk seeking behavioral choices when making collective choices between nests (Burns et al. 2016). The collective decision-making of the colony has also been shown to result in the avoidance of certain irrational choice behaviors observed in individual ants, including reducing the time to choose between potential nest sites (Sasaki et al. 2018, 2019). Recently, De Agrò and colleagues (2021) showed that ant foragers’ perceptions of food reward quality are based on the two reward options’ logarithmic values. Individuals were shown to be risk adverse when choosing between food rewards with the same mean values, preferring the fixed option, however this preference disappeared when ants were presented two logarithmically balanced alternatives (De Agrò et al. 2021).

In the current study, we characterized foragers’ learning and memories of aversive views when these views are associated with aversive, high effort outcomes, i.e. being kept within a brush-filled phial for set time periods. Forcing foragers into brush simulates areas along the homeward route that contain dense clutter, compelling the forager to struggle through in order to reach the nest with its food piece, increasing both their time and energy expenditure. Foragers’ behaviors were recorded using a trial-by-trial approach to describe navigational learning during natural tasks (Freas et al. 2019). We first studied the dynamics of view learning, as well as retention across non-reinforced trials. Second, we explored foragers’ perception of captivity duration by training foragers to associate sites along the route with two distinct fixed time periods within the brush (15s vs. 300s). Finally, we characterized foragers’ perception of risk when sites were associated with ‘Fixed’ or ‘Risky’ outcomes with the same mean duration across training (~150s). Here, the ‘Fixed’ outcome was associated with a constant period within the brush (150s) while the ‘Risky’ outcome was associated with a variable time period where foragers had a 50/50 chance on each trip of being held within the brush for either 1s or 300s. We found that *C. velox* foragers rapidly learn to associate the (previously positive) homeward route views with aversive outcomes, with as few as two prior experiences. These aversive view memories persist over multiple trips after the outcome is removed. Foragers were able to perceive differences in outcome severity, learning more rapidly and exhibiting more overall hesitations to views associated with more severe outcomes (300s) compared to less severe outcomes (15s). Finally, we show that foragers show significantly less apprehension to travel through sites associated with risky aversive outcomes compared to a fixed outcome with the same mean. The observed forager hesitation responses at these sites are in line with the perception of stimulus strength associated with their geometric averages.

## Methods

### Study site and species

Testing was conducted in June and July 2019 on a single *C. velox* nest located at an established field site ~6 km south of Seville, Spain (37°19′51′′N, 5°59′23′′W). *C. velox* inhabit visually cluttered semi-arid environments, densely covered in grass tussocks, scattered bushes and with distant stands of trees and man-made structures. While navigating, these foragers rely heavily on these visual cues to return to the nest and known food sites, creating stable routes between locations (Mangan and Webb 2012; Wystrach et al. 2015; Schwarz et al. 2017).

### Testing arena

A plastic square-shaped feeder (15cm × 15cm × 8cm) was sunk into the ground 12m from the nest entrance and was continuously stocked with crushed cookie pieces (Royal Dansk™). The smooth walls of the feeder prevented foragers that dropped in from exiting without being lifted out by the researcher. All vegetation in a 2m wide band from the nest to the feeder and in a 1m radius around both sites was removed using an edge trimmer. To entice foragers to collect food only from the feeder, an arena was erected using a 10cm high smooth plastic barrier, enclosing the nest and feeder site and restricting the nest to forage only within the arena. This arena was 1.5m in width and extended in a 75cm radius semi-circle around both feeder and nest (Fig. 1a). Two collection sites along the feeder-nest route were designated at 8m and 6m (Site 1 and Site 2 respectively) from the nest. To record inbound forager behaviour leading up to each site, two grids consisting of a 2 × 2 of 50cm squares were erected using string and metal pegs extending from each collection site 1m towards the feeder (ending at 9m for Site 1 and 7m for Site 2; Fig. 1a,b). Two sets of barriers were erected at 45deg angles creating a ~20cm gap at the edge of the grid to funnel foragers toward the centre of the arena’s width (Fig. 1a,b). To create two distinct panoramic scenes, at Site 1 the first set of erected barriers were 10cm high plastic walls identical to the walls of the arena, while the second set of barriers leading to Site 2 were 1.2m high (Fig. 1c). Additionally, to increase the panorama differences between sites, we placed a number of shorter 15–25cm visual landmarks consisting of stones and bricks around Site 2 (See Fig. 1c). This arena set-up was used for all three experiments.

**Fig. 1.**
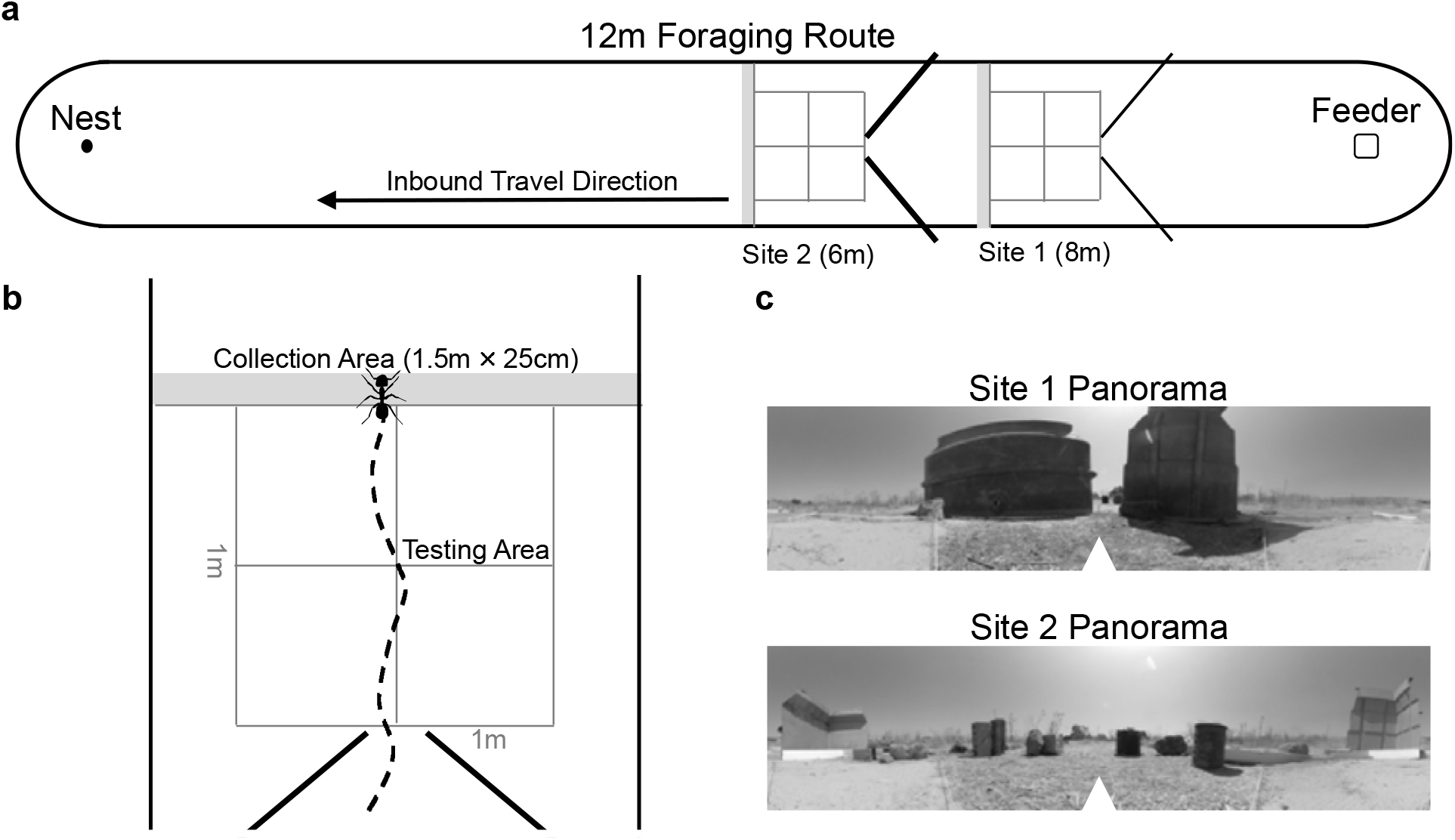
Diagram of the experimental set-up in all conditions. (a) Foragers were allowed to travel freely on the outbound trip to the feeder and collect food. After their release from the feeder, foragers travelled back to the nest through the testing areas at Site 1 and Site 2 and were collected based on the condition. (b) During collection conditions, foragers were collected within a 25cm (grey) area after passing 8m (Site 1) or 6m (Site 2) from the nest to allow the collector to remain as far back as possible before collection. After the allotted hold period, the collection phial was placed at the centre of the collection area and foragers were allowed to climb out and resume their homeward trip. The testing areas were arranged with blocking walls to both funnel foragers to the centre of the arena as they reached each testing area as well as to create distinct visual panoramas at each site. (c) Panoramic 360° photos of the surrounding visual cues at each collection site. In each photo, the arrow denotes the nest direction.

Upon the completion of the foraging arena construction, we allowed the nest two days to discover the feeder and begin consistently foraging. For these two days, any foragers reaching the feeder were allowed to enter and exit via a wooden ramp. At the onset of training this ramp was removed. As foraging began on the third day, it was expected foragers used in the experiments had some level of knowledge of the route before the onset of training and had learned the positive association between the views of the route and successful foraging trips. While the exact level of experience with the route prior to training may have individually varied, forager experience of the route during training was strictly controlled. When a researcher was not present to conduct training/testing, all foragers were restricted to a 20cm area around the nest using a plastic cylinder (~20cm in height).

### Procedure

#### Aversive learning tests

We initially tested foragers’ learning of aversive view memories by collecting inbound foragers as they reached Site 2 (Fig. 1a, Fig. 2). When approaching a view that has been associated with a negative outcome, foragers have been shown to hesitate leading up to the site, exhibiting bouts of scanning behaviour as well as attempting to avoid these sites via detours (Wystrach et al. 2020). In the current study, we collected two types of hesitation behaviour, scans and stops. Scans were defined as the ceasing of forager movement that was accompanied by the forager clearly turning on the spot, rotating in place. In contrast, stops were cataloged as the ceasing of forward movement with no accompanying rotation.

**Fig. 2.**
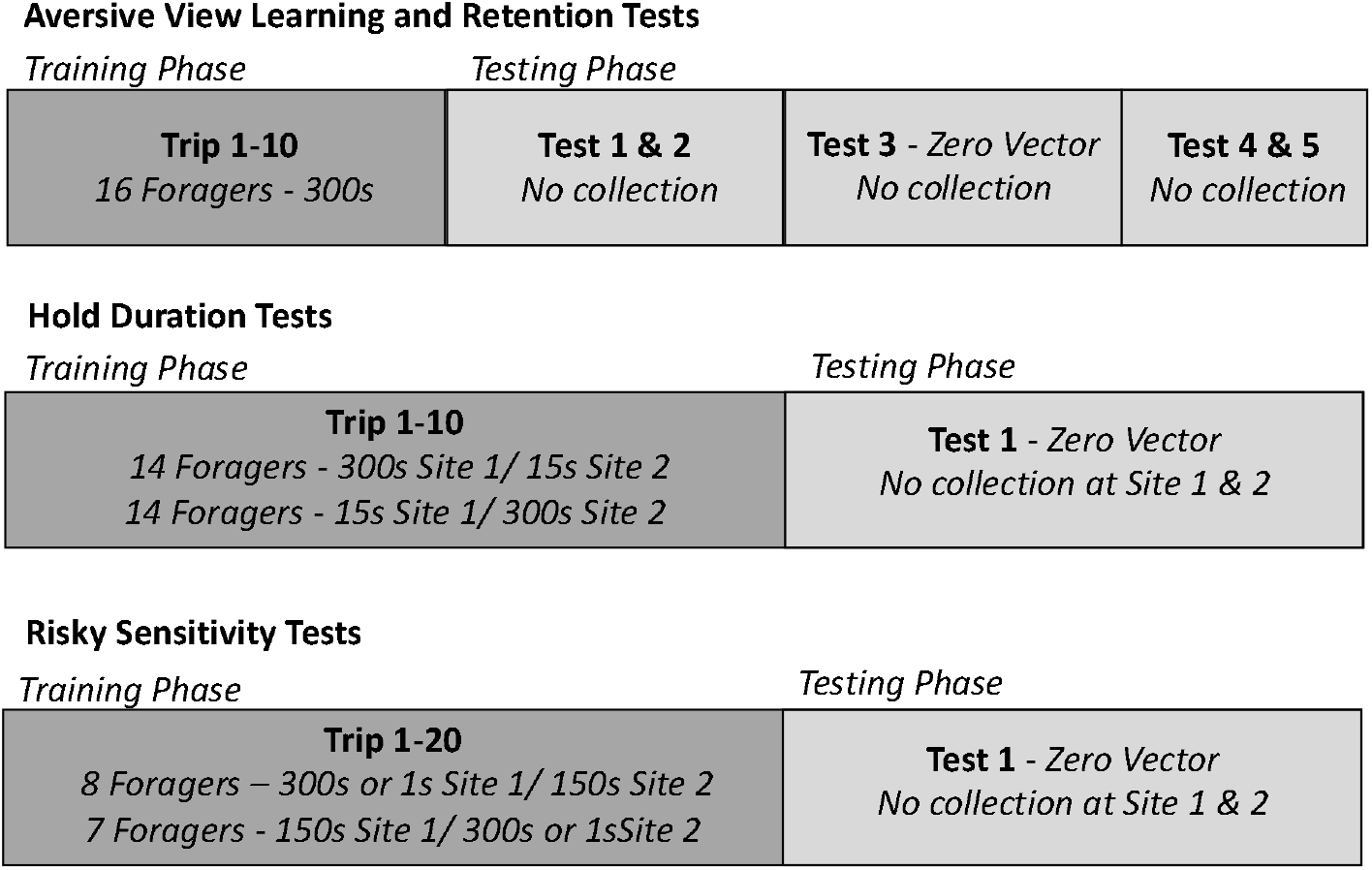
Timeline of training and testing procedures in all conditions.

During training, each forager (n = 16) was exposed to ten consecutive training trips, followed by five tests (Fig. 2). At the onset of training, foragers were allowed to travel freely from the nest to the feeder. Once in the feeder, foragers were individually marked using acrylic paint (Tamiya™), returned to the feeder and allowed to collect a cookie piece. After collecting their food piece, foragers were lifted out of the feeder by hand and allowed to travel through the arena to the nest. As foragers neared the grid at Site 2, their paths were recorded using an HD camera at 60 fps with a 3840 × 2160 pixels image size (GoPro™) positioned 1.2m above the grid facing down. Recording started just before the forager entered the grid area and ceased at collection. As the forager exited the grid (6m from the nest entrance), they were exposed to an aversive outcome in which they were held captive in a brush-filled phial. Specifically, foragers were collected within a 25cm area (grey area, Fig. 1a,b) using an opaque plastic 5cm diameter vial, which was filled with ~10cm of loosely packed grass brush from the surrounding vegetation, and held within this brush for a period of 300s. During this holding period, the vial was covered and placed in a semi-shaded area to prevent overheating. Additionally, the lid of this vial rested lightly upon the top of the brush, preventing foragers from standing on top of the brush during their hold period. After 300s, the vial was placed at the center of Site 2’s collection area (grey area, Fig. 1b) and tilted ~75° with the opening facing the nest to allow the forager to climb out of the brush and back onto the foraging route and resume navigating. This procedure was repeated for training Trips 2–10.

On the forager’s eleventh trip to the feeder (Test 1), individuals were released from the feeder and allowed to travel the full inbound route to the nest without being collected and held (i.e. in the absence of the aversive outcome). As in training, foragers were recorded at Site 2 beginning just before they entered the grid. As there was no collection during testing trips, recording ceased once foragers reached 25cm past the grid at the end of the collection area. All of these foragers reached the nest and freely entered. On their next foraging trip, foragers were tested twice. These inbound foragers were recorded and allowed to pass through Site 2 (Test 2), identically to the previous test, until they reached the nest entrance. As foragers reached within 20cm of the nest entrance (with their path integrator now near zero, termed Zero Vector, ZV) each forager was collected using an empty vial and returned to the foraging route 10m from the nest (Fig. 1a). These foragers were allowed to resume their nest-ward journey and were recorded while passing through Site 2 identically to previous tests (Test 3 ZV) without the corresponding vector state present during training at these sites. Returning foragers collected at the nest are described as zero-vector as their path integrator (PI) no longer provides directional information to the nest, however note that the PI system is constantly in use and foragers in this test are still accumulating PI information. Importantly the foragers’ PI states during zero vector testing do not align with their PI states at the sites while training. During foragers’ next two foraging trips to the feeder their homeward journeys at Site 2 were recorded (Test 4 and Test 5) identically to Test 1. After Test 5 all testing on the individual ceased and foragers were collected at the nest, marked as tested and then released.

### Hold duration tests

Next, we characterized whether foragers perceived differences in severity of aversive outcomes and responded differently to the associated views. Here, foragers (n = 14) were exposed to ten consecutive training trips where Site 1 was associated with a hold period of 300s while Site 2 was associated with a hold period of 15s (Fig. 2). A mirrored condition (Site 1 – 15s, Site 2 – 300s), was conducted on a second set of foragers (n = 14). Foragers were individually marked at the feeder and then released once they collected a food piece. During training (Trips 1–10), as foragers neared the grid at Site 1, they were recorded using the HD camera beginning just before the forager entered the grid area. As the forager exited the gird (8m from the nest entrance), recording ceased and they were collected within a 25cm area past the grid (grey area, Fig. 1a,b). Foragers were collected and held individually within the brush filled vial for 300s or 15 s (depending on condition) and then released back at the center of Site 1, using the procedure described in the previous experiment. After release, foragers were allowed to travel to Site 2 where they were again recorded within the grid at this site, then foragers were collected and held within the brush-filled vial upon exiting the grid (grey area, Fig. 1b). At Site 2, foragers were held for the other hold time before being released and then allowed to travel back to the nest with their food piece. On Trip 10, after release from Site 2, foragers were collected for testing as they reached the nest. These foragers were collected with no remaining vector (< 20cm from the nest) using an empty vial and immediately released along the route 10m from the nest (Fig. 2).

Released foragers were allowed to return to the nest and were recorded while passing through the grid at both Site 1 and Site 2 without the corresponding vector state present during training at these sites. After testing, foragers were collected, marked as completed and released at the nest.

### Risk sensitivity tests

In the final group of tests, we characterized foragers’ perceptions of fixed and risky aversive outcomes over 20 foraging trips. Here, for one set of foragers (n = 8), Site 1 was associated with a fixed aversive outcome, being held in brush for a period of 150s on every training trip, while Site 2 was associated with a variable outcome, with a 50/50 chance of being held for a longer (300s) or shorter (1s) period. Given the short period within the brush during the 1s hold time, special care was taken to confirm that this hold period did not start until foragers came in contact with the brush. A mirrored condition was conducted on a second group of foragers (n = 7) with these hold periods switched (Site 1 – 50/50 chance of a 300s or 1s hold period; Site 2 – 150s hold period). Foragers were individually marked as they reached the feeder and then allowed to collect food and return towards the nest. At Site 1, foragers were recorded as they entered the grid then collected and held identically to previous conditions. After the designated holding period, foragers were released and allowed to travel to Site 2 where they were recorded as they entered the grid and then collected and held. This training occurred for 20 trips. After the Site 2 release on Trip 20, foragers were tested by being collected with a zero-vector state as they reached the nest, released at 10m from the nest and their return trip through Site 1 and Site 2 was recorded without collection.

### Data digitization and analysis

Videos were digitized using GraphClick (Arizona Software). Paths were digitized by marking the ant’s mesosoma at 200ms intervals beginning when the forager entered the grid and ceasing once foragers were collected during training or when they reached 25cm past the grid edge during testing. Aversive view learning and memory were assessed by recording the number of hesitations exhibited by the forager leading up to collection. Two types of hesitation behavior were observed during testing, scans and stops and these were confirmed during video playback. We classified a ‘stop’ as a ceasing of forward movement with the ant remaining stationary until forward movement resumed. In contrast, ‘scans’ were classified as the ceasing of forward movement that is accompanied by the ant pirouetting, or rotating in place before resuming forward movement. Both of these behaviors were collected by the experimenter during the experiment and both positioning and behavior type was confirmed using video analysis. The quantity and position of both behaviors were recorded along the forager’s digitized paths. For statistical analysis, scan and stop behaviors were combined to create a total hesitation count for each training and test trip.

In the Aversive Learning tests, we compared hesitation numbers (Stops + Scans) across training/testing trips using a General Linear Model (GLM) for count data (Poisson loglinear) with Individual ants as a random effect. In the Hold Duration and Risk Perception tests, where foragers were collected for distinct hold periods at both Site 1 and Site 2, both *Site Number* and *Hold Condition* were analyzed as fixed effects. Post hoc pairwise comparisons of forager hesitations during the baseline during Trip 1 and after training/testing were conducted using p values corrected with the Bonferroni method. Within individual comparisons between hold regimes in the *Hold Condition* (15s v. 300s) and the *Risk Sensitivity* (Fixed vs. Risky) tests were compared using Wilcoxon Signed Rank Tests with p-values corrected with the Bonferroni method.

To further characterize the change in hesitation numbers at the Risky site, we calculated the change in hesitation (current hesitations minus hesitations on previous trip) number based on the outcome of the previous trip (held for 1s or 300s). For each individual forager, the mean hesitation change (excluding Trip 1) after a 300s hold time was compared to the mean hesitation change after a 1s hold time using a Wilcoxon Signed Rank Test.

For between test comparisons we chose to focus on forager hesitation numbers during training Trip 10 as, up to this point, the training schedule for each individual forager was consistent across all testing. Across testing, we compared the fixed and risky site hesitation numbers to those of the 15s site and 300s hold conditions using Mann-Whitney U tests. Finally, we further analyzed hesitations by calculating the predicted hesitation numbers along a logarithmic curve calibrated by the observed hesitations in the 15s, 150s (Fixed condition), and 300s conditions on Trip 10. We then compared the observed hesitations during Trip 10 of the Risky condition (μ = 3.33) to the predicted hesitations based on the conditions arithmetic (150.5s) and geometric (17.32s) hold times using one sample T-tests.

## Results

### Aversive learning tests

Foragers traveling through Site 2 at the onset of training (Trip 1) showed no signs of hesitation leading up to the collection site (μ ± *S.E.* = 0.0 ± 0.0; Fig. 3, 4) and this was used as the baseline for hesitation comparisons during training and testing. *Trip Number* had a significant effect on foragers’ hesitation numbers (*Z* = 3.84; p < 0.001). Post hoc comparisons showed that hesitations did not significantly increase from the baseline on training trips 2–4 (p > 0.05; Fig. 3, 4). Beginning on Trip 5 (*T* = 3.42, p = 0.01) and continuing through the rest of training (Trip 7– 10), hesitations were significantly higher than foragers’ baseline hesitation counts (p < 0.001). During testing (Test 1–5), post hoc comparisons revealed that hesitation numbers were significantly above baseline (Trip 1) during Test 1 (*T* = 6.39; p < 0.001), Test 2 (*T =* 3.19; p = 0.032), and Test 3 ZV (*T* = 3.26; p = 0.016; Fig. 3, 4). Beginning on Test 4 and continuing during Test 5, hesitation counts returned to baseline and were not significantly different from Trip 1 (p > 0.05). There was no significant difference between hesitation numbers during the last training trip, Trip 10 and Test 1 (*T =* 0.24; p = 1.00) as well as between Test 2 and Test 3 ZV (p = 0.87). Results suggest foragers showed significant signs of aversive view learning after three prior experiences and these hesitations increased throughout training, peaking at Test 1 (μ ± *S.E.*= 13.8 ± 2.9). Evidence of memory retention persisted for two trips after the aversive outcome was removed (Test 2 and Test 3 ZV) before hesitations returned to baseline (Test 4 and Test 5; Fig. 3). Finally, increased hesitations in zero vector ants confirm that the memories are associated, at least partially, with the views and not the forager’s vector state.

**Fig. 3.**
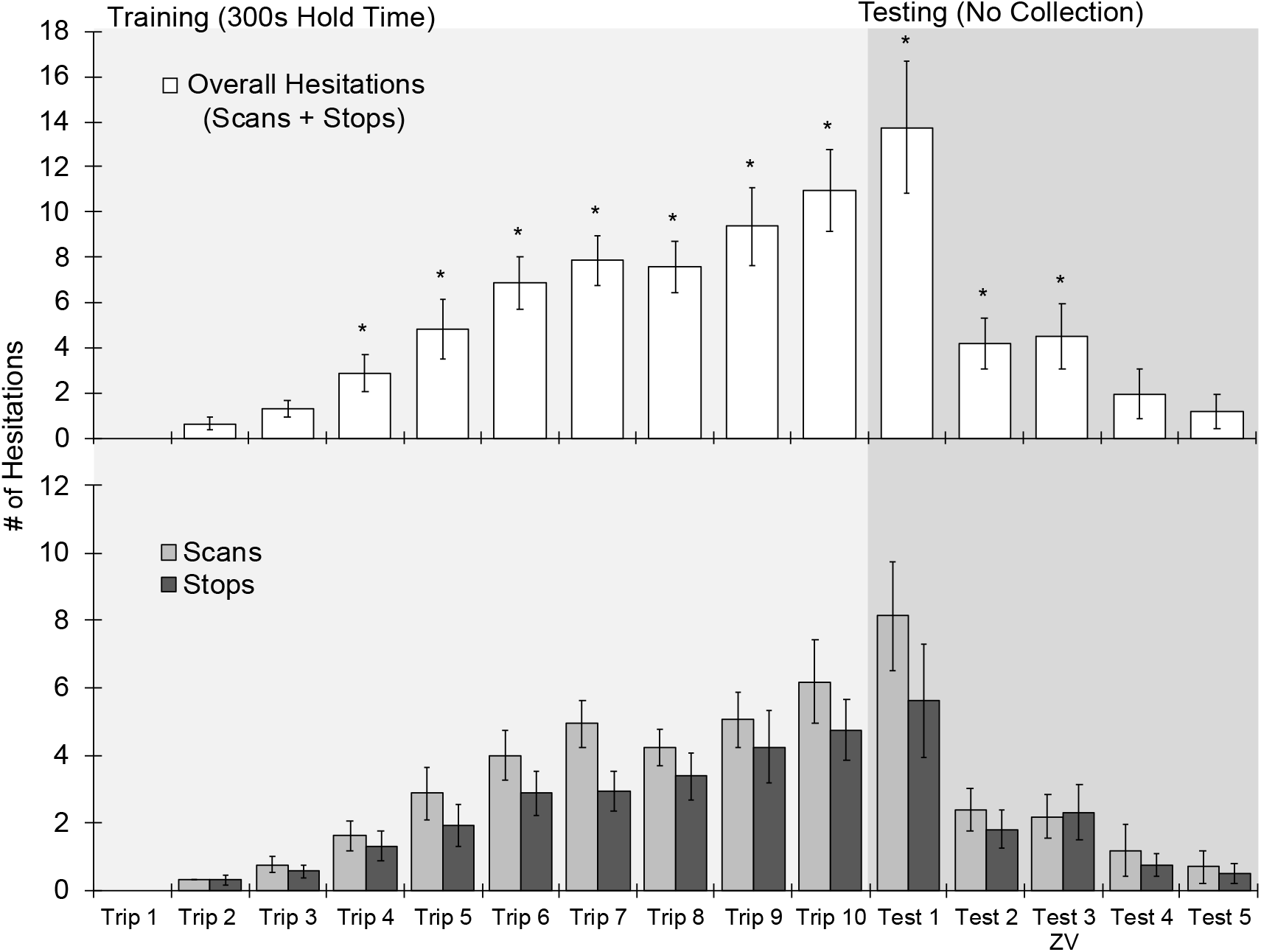
Forager hesitations during training and testing in the Aversive Learning tests. Foragers were collected at Site 2 during training (Trips 1–10), for 300s and were then allowed to return to the foraging route. During testing (Test 1–5), foragers were allowed to travel through Site 2 and return to the nest. (*a*) Mean hesitations across training and testing, consisting of scans and stops combined ± *SE. ‘*\*’ denotes training or test trips where forager hesitations are significantly above the Trip 1 baseline (p < 0.05). (*b*) Mean scans and stops across training and testing ± *SE*.

**Fig. 4.**
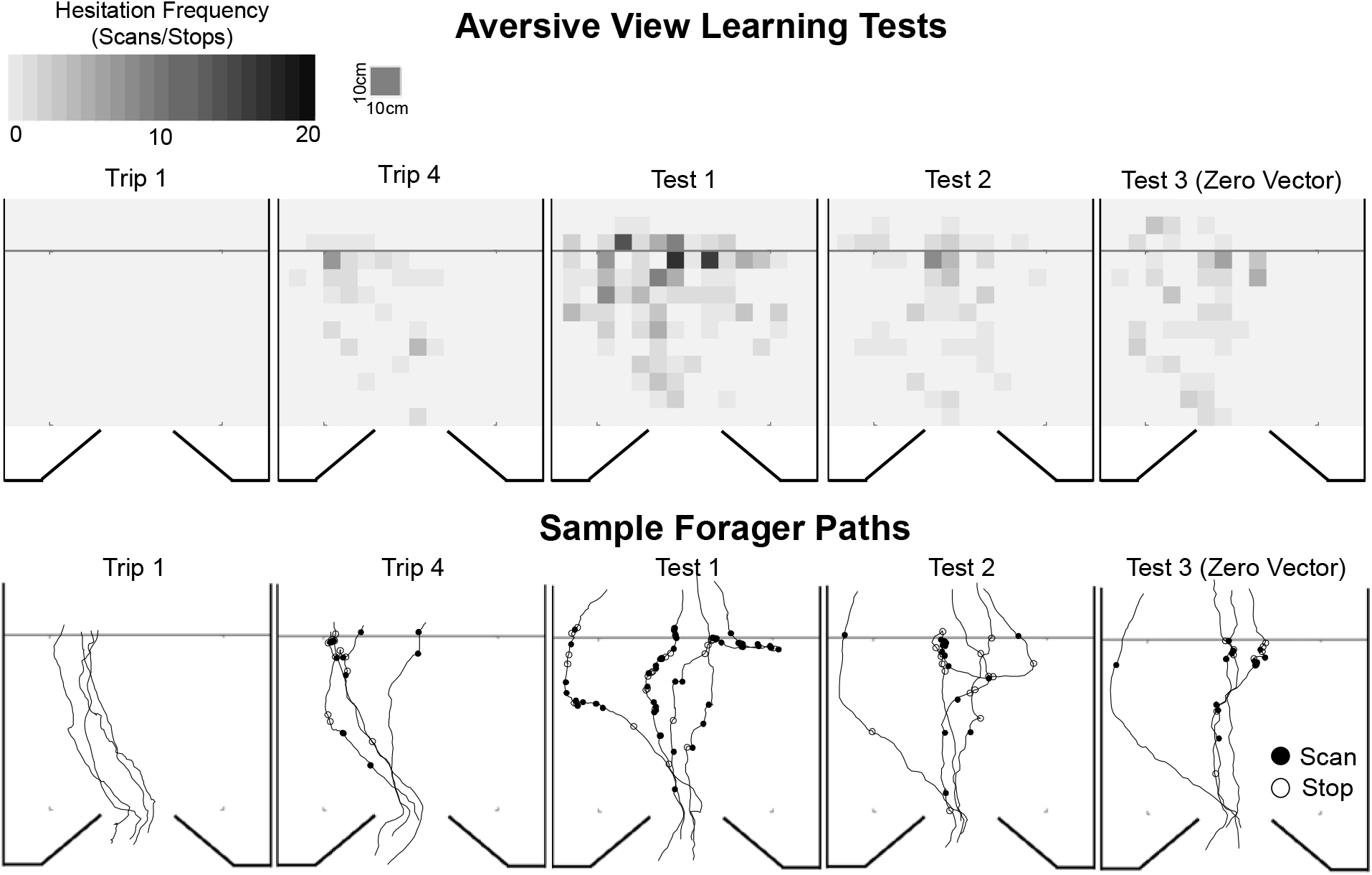
Heat maps of forager hesitation locations and sample paths in the Aversive Learning Tests. Heat maps of forager hesitations and five foragers’ sample paths during Trip 1, Trip 4, Test 1, Test 2 and Test 3. Closed black circles denote locations of forager Scans while open circles denote Stop locations. For heat maps of hesitations for all training and test trips in the Aversive Learning tests, see SFig. 1.

### Hold duration tests

Foragers at the onset of training (Trip 1) exhibited few pre-training hesitations leading up to both collection sites (15s Trip 1, μ ± *S.E.* = 0.89 ± 0.21; 300s Trip 1, μ ± *S.E.* = 0.64 ± 0.14; Fig. 5, 6) and these were used as the baselines for comparisons. Both *Trip Number* and *Hold Condition* had a significant effect on hesitations (*Trip Number*, *Z* = 6.21; p < 0.001; *Hold Condition*, *Z* = −3.47; p < 0.001) and there was a significant interaction between *Trip Number* and *Hold Condition* (*Z* =−4.98; p < 0.001). There was no significant effect of *Site Number* (condition mirroring) on forager hesitation numbers (*Z* = 0.34; p = 0.73) and *Site Number* showed no significant interaction with *Hold Condition* (*Z* = 1.05; p = 0.30) or *Trip Number* (*Z* = 0.85; p = 0.40).

**Fig. 5.**
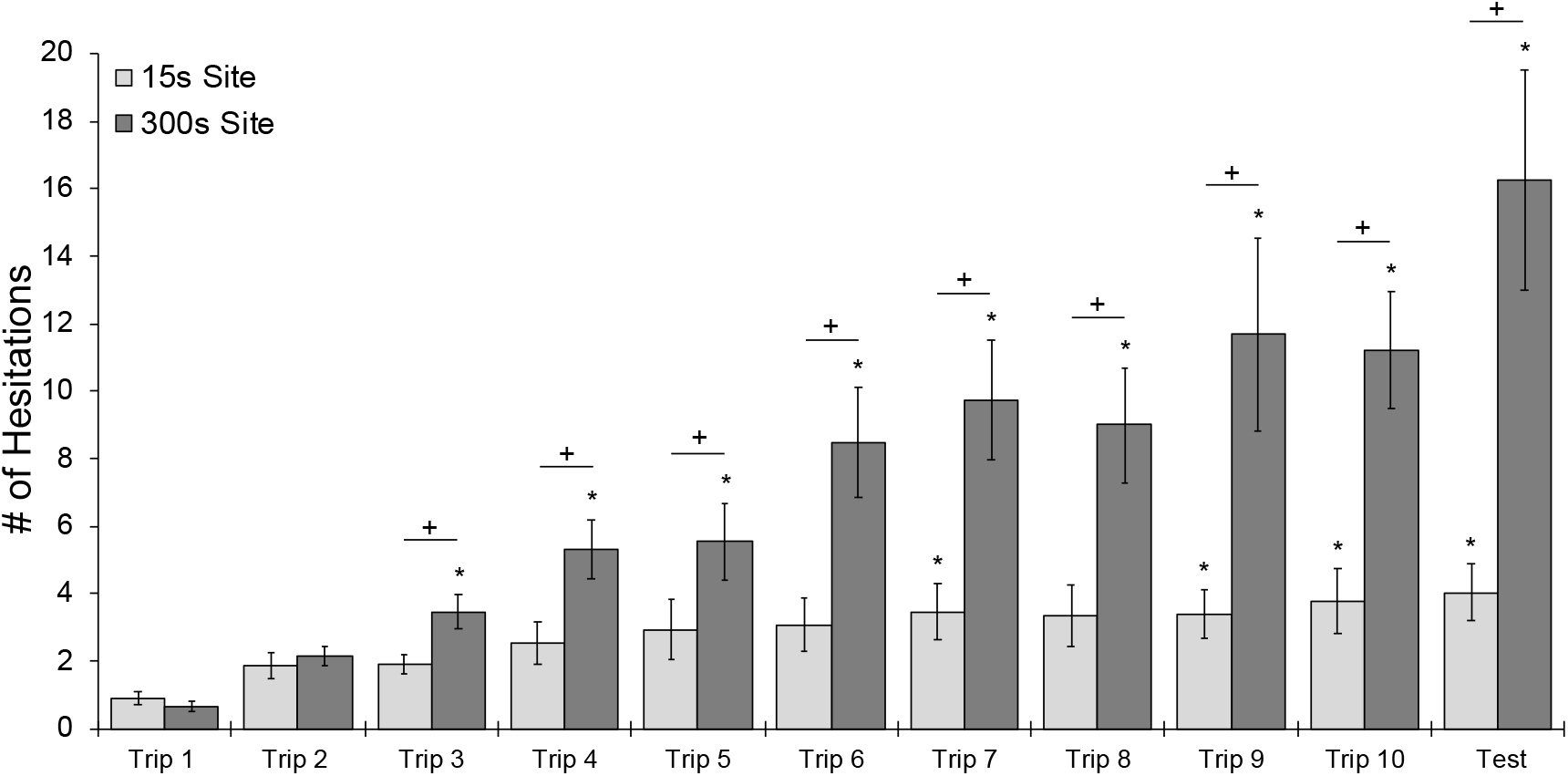
Forager hesitations during training Trips 1–10 and the zero vector test during Hold Duration tests. During training, foragers were collected at both Site 1 and Site 2 and held for either 15s or 300s before being released back at the site (hold periods for each site were randomly assigned at the onset of training for each individual). After training Trip 10, foragers were collected at the nest and tested as a ‘zero vector’ forager. During the test, foragers were allowed to travel through Site 1 and Site 2 without collection. Each *‘*\*’ denotes training or test trips where foragers’ hesitation numbers were significantly above the Trip 1 baseline (p < 0.05). Each ‘+’ denotes trips in which the forager showed significantly higher hesitation numbers leading up to the 300s collection site compared to the 15s site (p < 0.05).

**Fig. 6.**
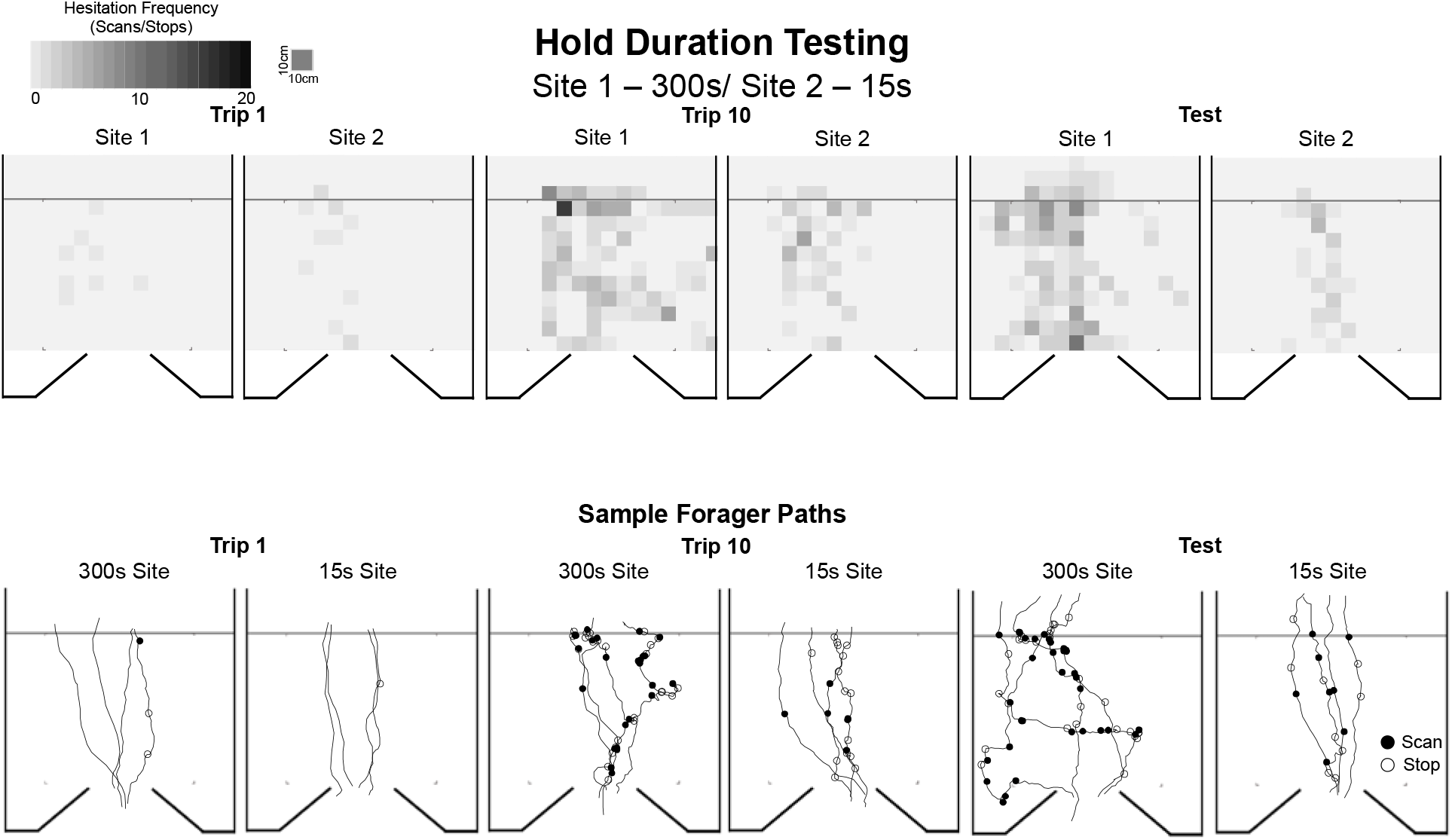
Heat maps of forager hesitation locations and sample paths in the Hold Duration testing. Closed black circles denote locations of forager Scans while open circles denote Stop locations. For heat maps of hesitations for all training and test trips in the Hold Duration tests, see SFig. 2,3.

At the 15s hold associated site, post hoc comparisons showed that hesitations did not significantly increase from the baseline on training Trips 2–6 (p > 0.05). Beginning on training Trip 7 (*T* = 3.11; p = 0.02) hesitations were significantly higher than foragers’ baseline hesitation counts, yet this significance increase disappeared during Trip 8 (*T* = 2.84, p = 0.06) before re-emerging for the final two training trips (Trip 9, *T* = 3.02; p = 0.04; Trip 10, *T* = 3.88; p = 0.01) as well as the zero vector Test (*T* = 3.69; p < 0.001). In contrast at the 300s hold site, post hoc comparisons showed that foragers learned the association after only two exposures, with hesitations significantly above Trip 1’s baseline beginning on Trip 3 (*T* = 2.38; p < 0.001). This significantly higher number of hesitations persisted through the rest of training on Trips 4–10 (p < 0.001) as well as the zero vector Test (p < 0.001). Comparisons between the final training trip, Trip 10, and the zero vector Test showed no significant difference in hesitations at either the 15s site (*T* = 0.19; p = 1.00) or 300s site (*T* = 1.14; p = 1.00).

Comparisons of forager hesitation numbers between the 15s and 300s sites showed that before training (Trip 1) foragers showed no significant differences between Site 1 and Site 2 (Wilcoxon signed-rank, Z = 43.5, p = 1.00). Foragers also showed no significant increase in their hesitations at the 300s site during Trip 2 (p = 0.23). Beginning on Trip 3 (p = 0.002), and continuing for the rest of training (Trips 4–10), foragers exhibited significantly higher hesitations at the 300s hold site compared to the 15s hold site (p < 0.005) and this difference persisted during the zero vector Test (p < 0.001). Foragers were able to perceive differences in outcome severity between two hold times, as foragers learned the association at the 300s hold site faster than the 15s site (Trip 3 vs. Trip 7) and showed higher hesitation counts associated with the more severe outcome associated site.

### Risk sensitivity tests

As in previous conditions, foragers at the onset of training (Trip 1) in both the Fixed and Risky conditions exhibited few hesitations leading up to the collection sites (Fixed Trip 1, μ ± *S.E.* = 0.60 ± 0.16; Risky Trip 1, μ ± *S.E.* = 0.69 ± 0.18; Fig. 7) and this was used as the baseline for future comparisons. Both *Trip Number* and *Hold Condition* (classified as Fixed or Risky) had a significant effect on hesitations (*Trip Number*, *Z* = 6.41; p < 0.001; *Hold Condition*, *Z* = 3.54; p < 0.001; Fig. 7) and there was no significant interaction between *Trip Number* and *Hold Condition* (*Z* = 1.1; p = 0.24). There was no significant effect of *Site Number* (condition mirroring) on forager hesitation numbers (*Z* = 0.47; p = 0.64) and *Site Number* showed no significant interaction with *Hold Condition* (*Z* = 0.34; p = 0.73) or *Trip Number* (*Z* = 0.91; p = 0.36).

**Fig. 7.**
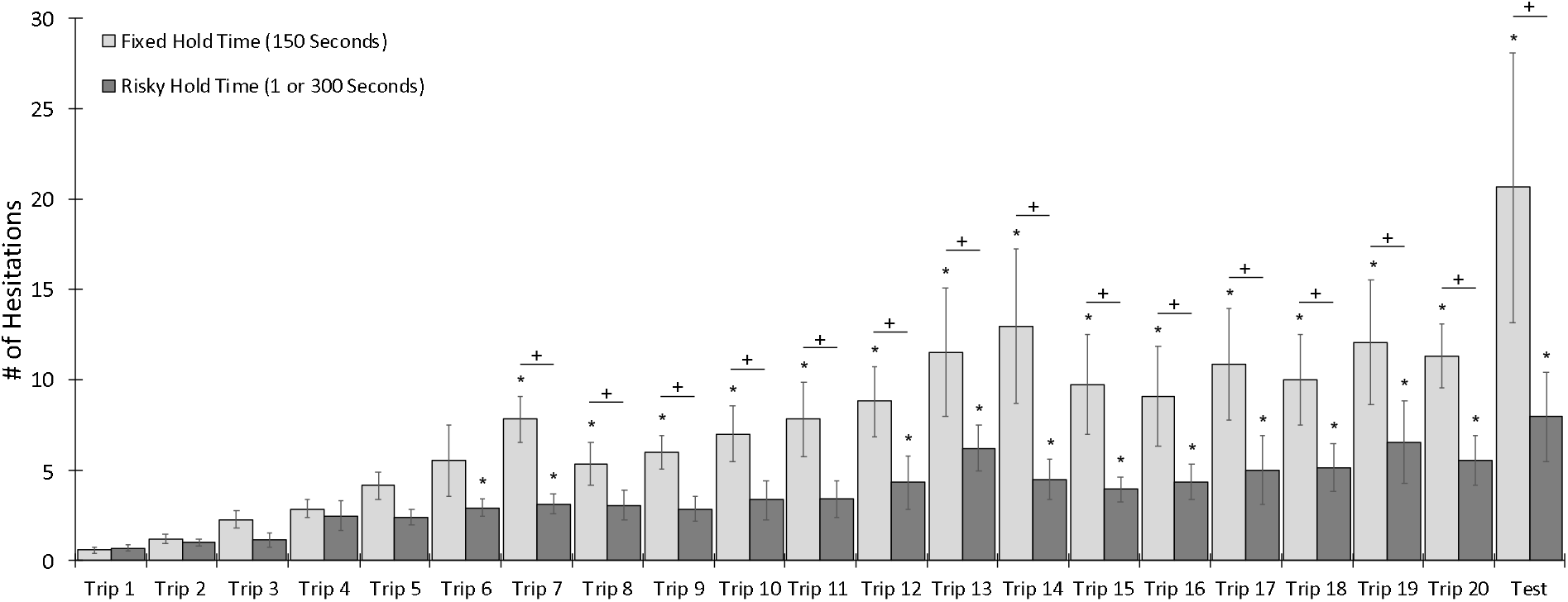
Forager hesitations during training Trips 1–20 and one zero vector test in the Risk sensitivity tests. During training, foragers were collected at both Site 1 and Site 2 and held within a brush filled container for either a fixed or risky period. At the fixed site, foragers were held for 150s, while at the risky site foragers had a 50/50 chance of being held for 1s or 300s. After the hold period, foragers were allowed to climb out of the container and return with their food to the nest. After training Trip 20, foragers were collected at the nest and tested as ‘zero vector’ foragers, by placing them back onto the route at 10m from the nest. During the test, released foragers were allowed to travel through Site 1 and Site 2 without collection. Each *‘*\*’ denotes training or test trips where forager hesitation numbers were significantly above the Trip 1 baseline (p < 0.05). Each ‘+’ denotes trips in which the forager showed significantly higher hesitation numbers leading up to the fixed (150s) collection site compared to the risky (1s/300s) site (p < 0.05). For heat maps of hesitations for all training and test trips in the Risky Perception tests, see SFig. 4,5.

At the Fixed site, post hoc comparisons showed that hesitations did not significantly increase from the baseline on training Trips 2–6 (p > 0.05; Fig. 7). On Trip 7, hesitations were significantly higher than foragers’ baseline hesitation counts (p < 0.001) and this significant hesitation increase persisted throughout the rest of training on Trips 8–20 (p < 0.05; Fig. 7) During the zero vector Test, forager hesitations were also significantly above baseline (p < 0.001).

At the Risky site, post hoc comparisons showed that hesitations did not significantly increase from the baseline on training Trips 2–5 (p > 0.05; Fig. 7). Hesitations were significantly above baseline during training Trip 6 (p = 0.02) and Trip 7 (p = 0.02), but not during Trip 8 and Trip 9 (p > 0.05). Beginning on Trip 10, hesitations were again significantly above baseline (p = 0.02) and this difference persisted through the rest of training (p < 0.05). During the Test, ZV forager hesitations were also significantly above baseline (p = 0.009; Fig. 7).

Comparisons of forager hesitations between fixed and risky hold schedules showed that, before training (Trip 1), foragers showed no significant differences between Site 1 and Site 2 (Wilcoxon signed-rank; Z = 10.50; p = 1.00). During training, foragers also showed no significant difference in their hesitations between sites during Trip 2 (Wilcoxon signed-rank; Z = 32.5; p = 1.00) and this persisted through Trip 6 (Wilcoxon signed-rank; Z = 11.02; p = 0.20). Beginning on Trip 7 (Wilcoxon signed-rank; Z = 5.01; p = 0.004) and continuing through the rest of training (Trips 8–20), foragers exhibited significantly higher hesitations at the Fixed hold site compared to the Risky hold site (p < 0.05) and this difference was also present during the zero vector Test (p < 0.001).

During training at the Risky site, changes in hesitation numbers suggest that the effect of training was continuously regulating hesitation behavior up and down based upon differences in the expected outcome and forager’s experience on each trip (Fig. 8a). After experiencing the highly aversive 300s outcome, hesitations are regulated upward (mean hesitation change ± *S.E.* = +1.13 ± 0.44) while experiencing the less aversive 1s outcome resulted in hesitations being regulated downward (mean hesitation change ± *S.E.* = −0.46 ± 0.28) and these changes based on the last experience were significant (Wilcoxon signed-rank; Z = 18.00; p = 0.02).

**Fig. 8.**
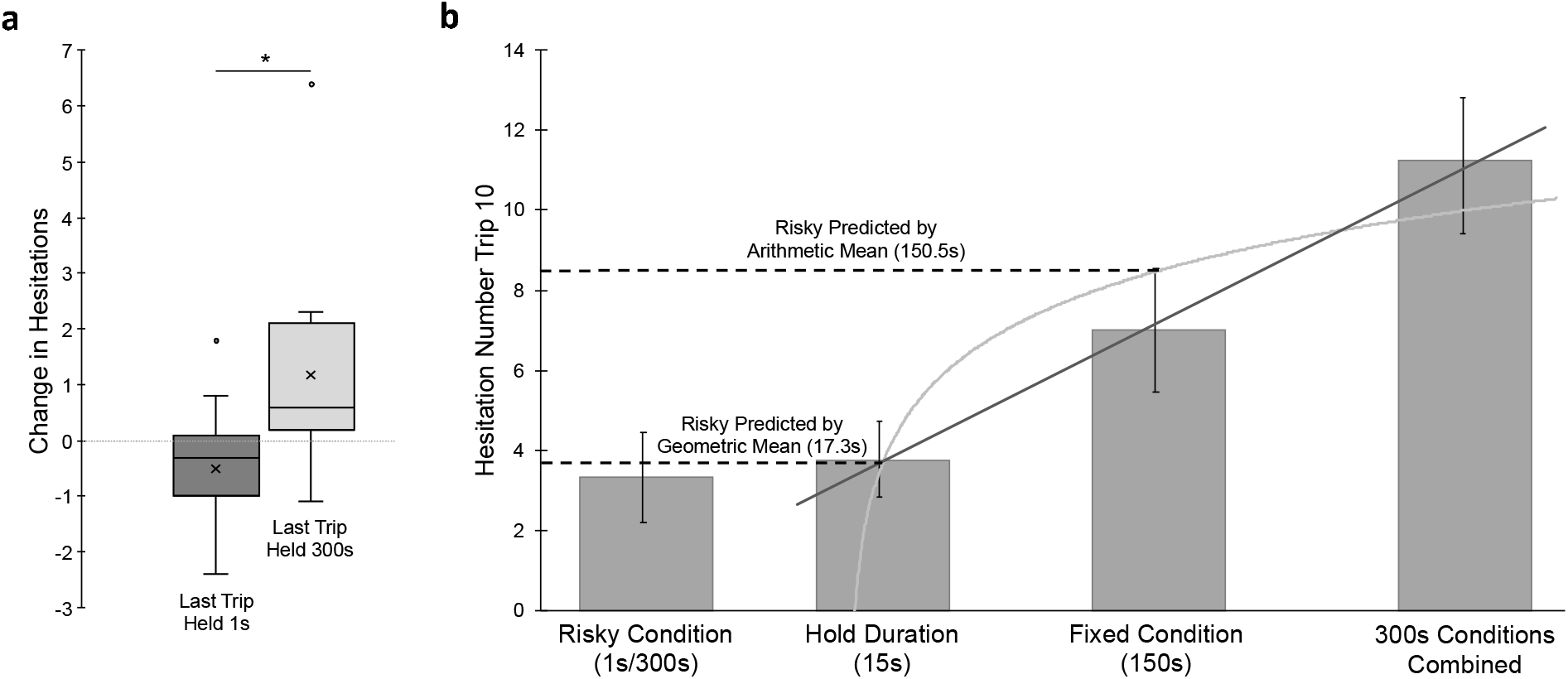
(a) Mean Change in hesitation numbers in the Risky condition across training based on the outcome of the previous trip (Held 300s or 1s). Mean change across training was calculated per individual. Box plots shows the median hesitation change (middle line), mean hesitation change (⍰) and 25th and 75th percentile (box) while the whiskers extend to min and max values (excluding outliers). Outliers were defined as values 150% of the IQR beyond 25th and 75th percentile and represented as individual points. * denotes a significant difference between conditions (p < 0.05). (b) Predicted hesitations at training Trip 10 along a logarithmic curve (grey line) calibrated by hesitation numbers in the Hold Duration (15s), Fixed condition (150s), and the combined 300s conditions (mean hesitations ± *S.E.*). The Risky condition’s mean number of hesitations on Trip 10 is plotted on the left, with the predicted value of either the condition’s geometric mean hold time of 17.3s or its arithmetic mean hold time of 150.5s. The observed hesitations during Trip 10 of the Risky condition (μ ± *S.E* = 3.3 ± 1.1) fell well below the predicted hesitation number given the arithmetic mean hold time (3.33 observed vs. 8.5 predicted) yet the logarithmic curve of predicted hesitations (3.8) falls within the standard error of the observed hesitations at the condition’s geometric mean. The observed lower hesitations during risky training aligns with the principle that the forager’s perception, and resulting hesitation behavior, of the aversive outcome has a logarithmic relationship with stimulus strength (y = 2.1824 × ln(x) − 2.4646).

### Between Condition Comparisons

During Trip 10, forager hesitations in the two 300s conditions (the Aversive Learning tests and Hold Duration tests) showed no significant differences (Mann-Whitney U; U = 210.5; p = 0.37) and these data were combined for future comparisons. Hesitation numbers during the Fixed 150s condition were significantly lower than the 300s conditions (Mann-Whitney U; U = 196.5; p = 0.048) and significantly higher than the 15s condition (Mann-Whitney U; U = 196.5; p = 0.002). In contrast, forager hesitations in the Risky condition, (mean hold time = 150.5s) were not significantly higher from the hesitation numbers of foragers in the 15s condition of the Hold Duration tests (Mann-Whitney U; U = 196.5; p = 0.741) despite the order of magnitude difference in mean hold time (150.5s vs. 15s; Fig. 8b).

Finally, observed hesitations during Trip 10 of the Risky condition (μ = 3.33) were compared along the logarithmic curve of predicted hesitation numbers (y = 2.2076 × ln(x) − 2.544; R = 0.9293) at both its arithmetic (150.5s) and geometric (17.32s) mean hold times. At Trip 10, observed hesitations in the Risky condition were not significantly different from the predicted hesitation number (3.75) based upon the geometric mean hold time of 17.32s (One sample T-test; *T* = 0.377; p = 0.712; Fig. 8b). In contrast, these observed hesitations significantly differed from the predicted hesitation number (8.53) based upon the arithmetic mean hold time of 150.5s (One sample T-test; *T* = 4.67; p < 0.001).

## Discussion

In all tests, foragers traveling through the grid leading to their first collection exhibited either no or minimal hesitation behaviours. Hesitations, including scanning behaviours, typically occur when there are increases in navigational uncertainty due to inexperience, cue conflicts or decreases in view familiarity (Wystrach et al. 2014). Given the high likelihood that foragers had multiple experiences of the route during previous successful foraging trips to the feeder, it was expected that hesitation numbers during Trip 1 would be low. When foragers were trained at either site along the homeward route with a captivity period in the brush filled container, these individuals learned an association between the views preceding the collection site and the outcome, showing a significant increase in hesitation numbers when encountering these views on the following trips. Foragers retained these view-based associations even after the outcome was removed for multiple trips. Trained foragers continued to hesitate above their baseline on the next three trips where they were allowed to pass through without collection, suggesting the behavioral response was not based solely on the previous foraging trip. Foragers tested without a corresponding vector state showed no change in hesitation numbers, meaning the association was tied primarily to the view memory and not an association between the vector state and outcome.

### Aversive view learning

Models of visual navigation currently rely solely on the positive valence, or attractiveness, of familiar views which inhibit search behaviour (turning) and induce forward movement (Wystrach et al. 2011b; Baddeley et al. 2012; Ardin et al. 2016; Kodzhabashev and Mangan 2015). These positive valance memories involve reinforcement learning of the associated inbound views that lead to the nest, likely reinforced upon the forager’s arrival, though the exact reinforcer remains unknown. Yet, recent work has demonstrated that views can also be associated with negative outcomes leading these views to develop a negative valence, or aversiveness, which inhibits forward movement and induces hesitations, turns and scanning behaviour (Wystrach et al. 2020). Such behaviours increase the likelihood that foragers may avoid the negative outcome experienced on the old route and return to the nest quickly along a new route, leading to these new views developing a positive valance and the formation of detours (Le Möel and Wystrach 2020, Murray et al. 2020; Wystrach et al. 2020).

In the current study, just as in Wystrach et al. (2020), our results show rapid acquisition of aversive view memories at specific spatial locations associated with negative outcomes, resulting in increased hesitations leading up to these sites. Views that previously had a positive association, formed during the initial formation of the homeward route, subsequently become negative when associated with the experience of struggling within the brush. Unlike previous work, here foragers were unable to form new positively reinforced routes detouring around these negative outcomes as all available homeward routes resulted in collection. Once the negative outcome was removed, foragers were shown to take two exposures to the route to re-learn its positive association, reinforced through re-entering the nest (during Test 1 and Test 3 ZV), and reduce hesitations to baseline. As Test 2 did not result in the forager successfully entering the nest (due to collection for Test 3 ZV testing), it is likely the observed lack of a decrease in hesitations between Test 2 and Test 3 (Fig. 3) was influenced by the missing reinforcement of re-entering to the nest, rather than reaching a zero-vector state or experiencing the nest panorama. This result hints that positive reinforcement of the route views upon a successful foraging trip may trigger only once the forager enters the nest.

### Outcome severity and risk perception

In associating views with negative outcomes, foragers are able to distinguish between levels of severity of outcome, which was evident both in the acquisition rates and overall hesitation responses. Foragers in the Hold Duration tests rapidly learned the association at the 300s site, showing increased hesitation behaviour after only two previous exposures. In contrast, these same foragers required six exposures to show increased hesitation behavior at the 15s site. Overall hesitations also differed by outcome severity, with foragers exhibiting significantly more hesitations associated with the 300s site (μ = 11.7) than the 15s site (μ = 3.8) during the final training trip (Trip 10). In contrast, foragers trained in the Risk Perception tests at the fixed 150s site, exhibited a hesitation response mid-point between these two extremes (μ = 7.3). These differences suggest foragers were able to perceive differences in time spent struggling within the brush and recalled this outcome severity on subsequent foraging trips, leading to distinct levels of aversion behavior expressed at each site.

Risk variance did not affect how quickly foragers learned the negative association, but had a significant effect on the degree of hesitation that developed. Over the final five training trips, foragers exhibited twice the number of hesitations at sites that resulted in a fixed negative outcome (150s; μ = 10.7) compared to sites leading to risky outcomes (1s or 300s; μ = 5.3) suggesting foragers perceived the risky outcome as less severe than the fixed outcome despite these schedules having the same mean hold time over the course of training (~150s). This increased hesitation number at the fixed site versus the risky site persisted during the test (μ = 20.6 and 7.9 respectively). Furthermore, hesitation number at the risky site was shown to increase or decrease based upon the outcome experienced on the forager’s previous trip. When foragers experienced a severe 300s hold time on the preceding trip, hesitations increased (μ = 1.16) while when the less aversive 1s hold time was experienced on the preceding trip, hesitations decreased (μ = −0.51). These changes indicate that forager’s hesitancy to pass through the site is being continuously regulated up or down with each new experience of the site. During each training trip, the hesitation behavior expressed represents the forager’s level of aversion to the expected outcome while each new experience regulates this expectation.

Additionally, the hesitation data presented here shows that foragers’ perception of risky aversive outcomes was not optimal in terms of the true value of the outcome. Many studies of preference variance in reward quality/quantity show a general risk aversion tied to the animal’s perception of rewards balanced only by mean (Kacelnik and Bateson 1996). This has been demonstrated in ants, as De Agrò et al. (2021) showed that ant foragers were risk averse when reward options were balance by mean value. Yet, fixed option preference disappeared when the two reward options were altered to be geometrically balanced (De Agrò et al. 2021). This finding makes sense if animals perceive reward value on a logarithmic scale. For positive outcomes, when the geometric average of a risky reward option falls below the fixed reward option, animals should perceive the fixed option as preferable. Our research suggests that this relationship also fits with the study of variability of negative outcomes. When the geometric average of a negative outcome falls below the fixed outcome, animals should perceive this risky outcome as less severe and become risk seeking, which aligns with our results. In the Risk Sensitivity tests, the two outcomes were only balanced by true value (150s vs. 150.5s), while the geometrical average of the hold duration of the Risky site 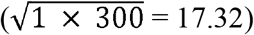 was lower than the Fixed site 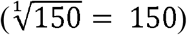. This corresponded with an overall lower level of hesitations at the risky outcome site compared to the fixed outcome site. While we did not test purely geometrically balanced Risky vs. Fixed outcomes, the perceived stimulus strength of the Risky site based on the geometrical average would make it very similar to that of the 15s outcome in the Hold Duration tests (17.32s vs. 15s). Interestingly, the hesitation levels during training (Trip 10) between the 15s outcome (μ = 3.8) and the Risky outcome (μ = 3.3) were not significantly different from one another despite the order of magnitude difference in actual mean hold duration (150.5s vs. 15s; Fig. 8b). Additionally, we calculated a logarithmic curve of expected hesitations based on those observed during the 15s, 150s and 300s conditions on Trip 10 (Fig. 8b). When hesitations observed during Trip 10 at the Risky site were compared with this curve, observed and predicted hesitations did not significantly differ at the geometric mean (17.32s) but did significantly differ from the predicted hesitations at the Risky condition’s arithmetical mean (150.5s), demonstrating that the geometric mean hypothesis should be favored over the arithmetic mean. This provides further evidence that there is a logarithmic relationship between captivity duration and forager’s response, suggesting foragers are perceiving the outcome severity logarithmically rather than its true value.

Stimulus strength perception is typically confirmed by animal’s choices of varying rewards and is used to explain why animals are typically risk averse to variable rewards (see Scalar Utility Theory, Kacelnik and El Mouden 2013). The current results indicate that such factors also predict risk seeking behavior to negative outcomes. Here, the foragers faced with two identical mean hold times perceived the variable outcome as less severe, and equal to a hold time almost an order of magnitude lower than its true value (17.3s vs. 150.5s) and thus respond less negatively to the associated views, on par with hesitations to a fixed 15s hold time (Fig. 8b). In our Risk Perception testing, the constant site always results in a 150s hold while the risky site may result in a 1s or 300s hold time. Along a logarithmic curve, this 150s hold time would be perceived by the forager as 150 times worse than the 1s hold time while 300s is only two times worse than 150s. If these outcomes were balanced by geometric mean rather than true mean, for example altering the hold times to a constant 15s hold time and a risky schedule of a 50% chance of either 1s or 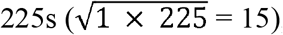, we would expect the forager’s perception of the aversive outcomes to be equal and result in identical hesitation numbers. While the current findings clearly point to the logarithmic relationship between the outcome’s severity and the forager’s aversive response, future work could help further untangle the ant’s perception of aversive outcomes, effort and risk by testing truly geometrically balanced fixed and variable outcomes.

### Underlying neural mechanisms

Specific view memories must be stored through some change within the brain (memory trace) along with their current corresponding positive or negative valence, based on previous experiences along the foraging route. Each memory trace must also be able to be altered between attractive and aversive valences and result in changes of forager behaviour when these views are next experienced.

The use of learnt route memory in ants involves the Mushroom bodies, MB (Büehlmann et al. 2020; Khamil et al. 2021), and its known neural circuitry can explain the storage and recall of visual as well as olfactory memories (Heisenberg 2003; Ardin et al. 2016; Webb and Wystrach 2016; Wystrach et al. 2020). Visual information enters the MB via projection neurons from the optic lobes (Habenstein et al. 2020). An individual view can be represented neurally within the MB through activation patterns of Kenyon Cells (KC), which project onto a number of motor output neurons (MBON). Each MBON conveys an attractive or aversive valence, and changes in synaptic strength between KCs and MBONs by activation of dopaminergic neuron mediating negative or positive experiences, modulates the association between a stimulus and its outputted valence (Cohn et al. 2015; Aso and Rubin 2016). Here, changes within these synaptic compartments mediate the view’s current overall valence by weighting the attractive and aversive valences of the forager’s experiences during previous trips to control the forager’s steering behavior. More specifically, the aversive outcome of being captured must result in dopaminergic neurons decreasing the connection strength between the recently encountered view specific pattern of Kenyon cells and attractive valence MBONs (Wystrach et al. 2020). This triggers the hesitation behaviours observed leading up to the collection site after training foragers with the aversive outcome.

The current study findings support our current understanding of the memory dynamics within the circuity of the MBs, mostly stemming from olfactory memory work in other insects. First, the aversive response increased across repeated trial until reaching a plateau (Fig. 7), as observed in drosophila olfactory conditioning (Beck et al. 2000). Second, while aversively trained foragers were highly hesitant to travel through these sites, no forager refused to cross the line despite the aversive association, even when they lacked a corresponding vector (Fig. 4; Fig. 5), suggesting that both attractive and aversive valence memory traces are simultaneously at play. Thus, it is likely some underlying attractive valence associated with these views persists in different MBONs, despite the acquired association with the aversive outcome, as demonstrated in fly’s MB (Boto and Ramaswami 2021). Third, the changes in hesitation number in the risky condition indicate that forager behaviour is being continuously regulated based by each new trip’s experienced outcome (Fig. 8a). Thus, current experience continuously regulates the forager’s expected outcome at the site, as observed in the fly’s MB (Cohn et al. 2020). However, the persistence of hesitations after the aversive outcome was removed show that valence persist, and is thus the result from an accumulation of experiences over multiple previous trips, not the last experience alone. Finally, the reduction in hesitation following a 1s hold time in the risky condition (Fig. 7) shows that this experience led to a net gain in positive valence, even though the ant has been captured. This supports the idea that learning in the MB follows a prediction-error rule (Bennett et al. 2021). Learning is dependent on the discrepancy between the current experience and the expected one (Rescorla and Wagner 1972). In other words, being captured for 1s at a site where one has been previously captured 300s mediates a positive reinforcement, leading to a decrease in aversion. Overall, these results indicate that various dopaminergic neurons are continuously modulating connection strength of various aversive and attractive valence MBONs based on the difference between the expected outcome and the experienced outcome on that trip.

## Conclusions

We found that *C. velox* foragers rapidly learn to associate views with aversive outcomes, showing increased hesitations at these sites, in some cases after only two previous experiences. Such memories are not solely based on the forager’s most recent trip, as individuals continued to showed increased hesitation at these sites after the aversive outcome was removed, suggesting these aversive memories persist over multiple trips. Foragers were also able to perceive differences in outcome severity, learning more rapidly and exhibiting more hesitations at a site associated with a severe outcome (300s) when compared to a less severe outcome (15s). Additionally, we show that the foragers show significantly less apprehension to travel through a site associated with a risky aversive outcome compared to a fixed outcome with the same mean and that forager hesitation responses at these sites across experiments was in line with the logarithmic relationship between stimulus strength and perception. Finally, our findings fit within the current modeling of view-based route learning and memory in the mushroom bodies of the insect brain. The behavioral dynamics observed here align well with the complex and parallel memory dynamics of the MB as studied in the context of olfaction in flies. In closing, a final intriguing question remains. Namely, the foragers’ response to different hold durations suggests that they can somehow quantify or estimate their duration of capture, yet the mechanism by which this duration estimate in accomplished currently remains unknown.

## Funding

This work was supported by a Mitacs Globalink Research Award (PI: CAF) and Natural Sciences and Engineering Research Council of Canada Discovery grants (#04133 and #2020-03933).

## Competing Interests

The authors declare no competing or financial interests.

## Ethical standards

There are no ethical requirements for working with insects in Spain. Manipulations were non-invasive and all individuals were returned to the nest after testing.

## Notes

### Competing Interest Statement

The authors have declared no competing interest.

