## Supplemental Figures for "Aversive View Memory and Navigational Risk Sensitivity in the Desert Ant, *Cataglyphis Velox*"

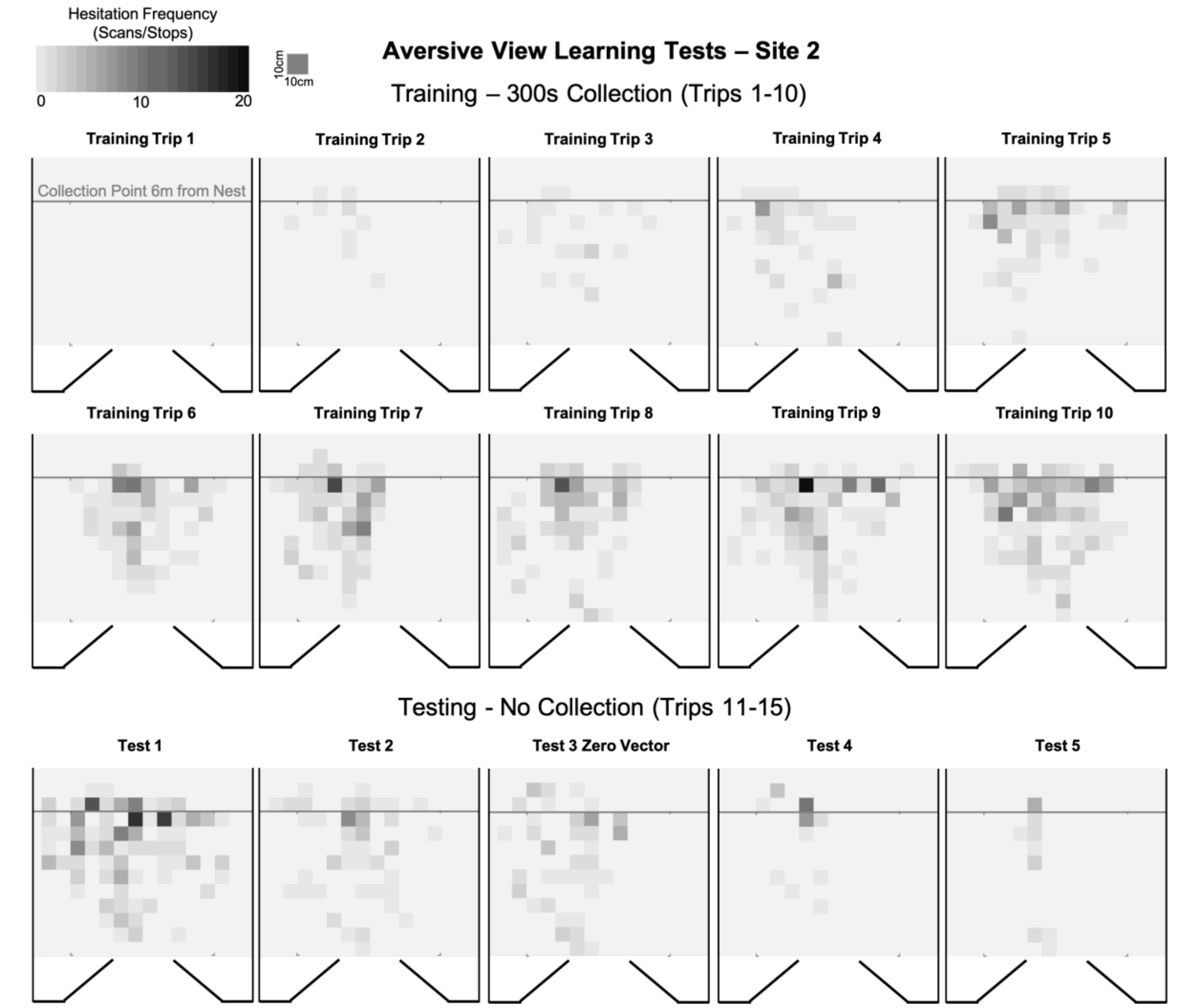


**SFig. 1.** Heat maps of forager hesitation locations at Site 2 in the Aversive Learning Tests. During training Trips 1–10, foragers (n = 14) were collected and held for 300s after they exited the testing grid (grey line, Collection Point) 6m from the nest entrance. During Test 1–5, foragers were not collected and instead allowed to travel through Site 2 and to the nest entrance. Foragers were tested with no corresponding homeward vector during Test 3 (Zero Vector).

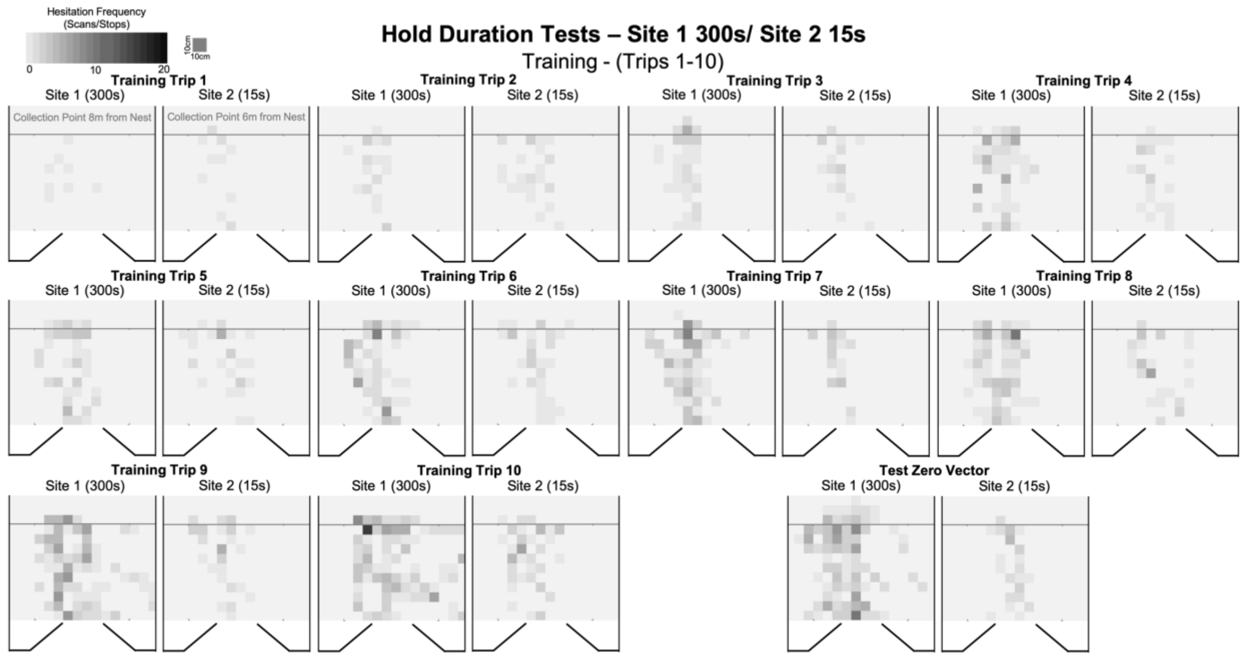


**SFig. 2.** Heat maps of forager hesitation locations in the Hold Duration Tests (Site 1 – 300s / Site 2 – 15s). Here, Site 1 was associated with a 300s hold time and Site 2 with a 15s hold time. During training Trips 1–10, foragers (n = 14) were collected after they exited the testing grid (grey line, Collection Point) at both Site 1 and 2 (8m and 6m from the nest entrance respectively). During the Test Trip, foragers were tested with no corresponding homeward vector (Zero Vector) and allowed to pass through both sites and travel to the nest entrance.

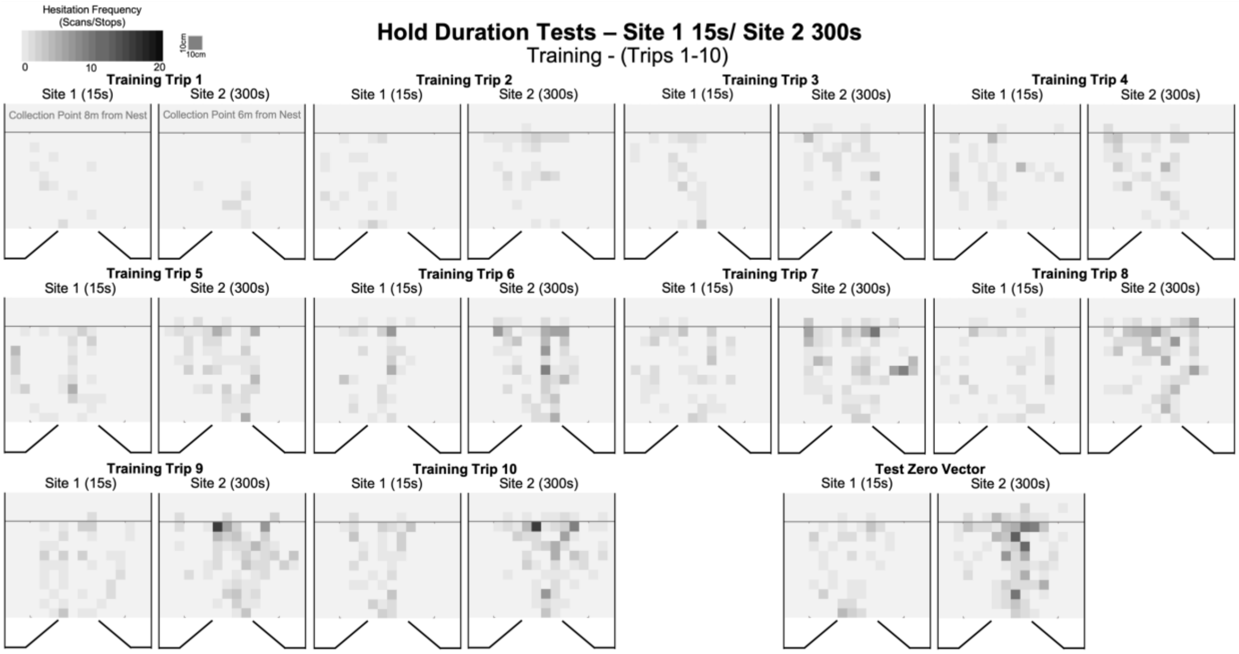


**SFig. 3.** Heat maps of forager hesitation locations in the Hold Duration Tests (Site 1 – 15s / Site 2 – 300s). Here, Site 1 was associated with a 15s hold time and Site 2 with a 300s hold time. During training Trips 1–10, foragers (n = 14) were collected after they exited the testing grid (grey line, Collection Point) at both Site 1 and 2 (8m and 6m from the nest entrance respectively). During the Test Trip, foragers were tested with no corresponding homeward vector (Zero Vector) and allowed to pass through both sites and travel to the nest entrance.

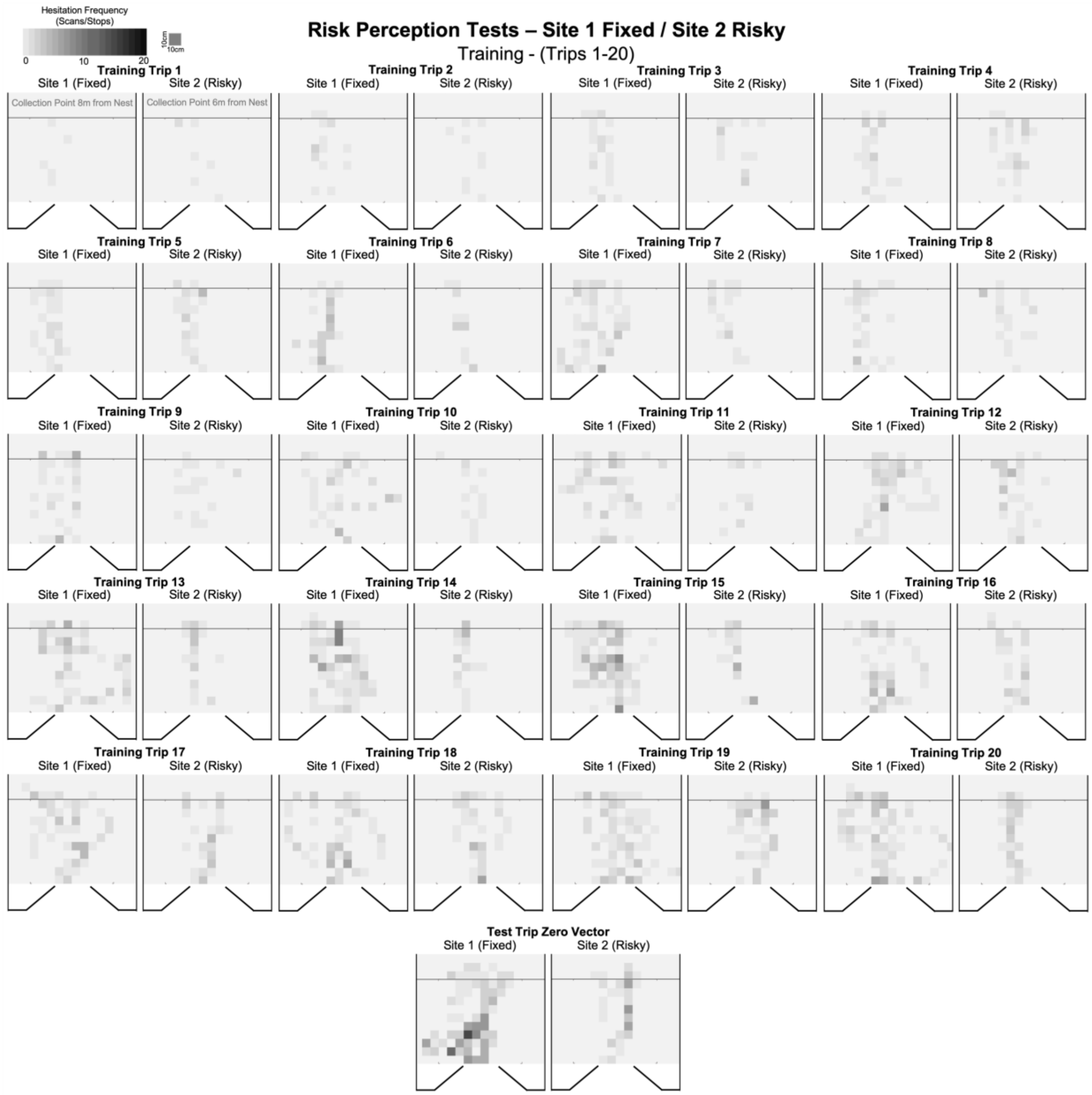


**SFig. 4.** Heat maps of forager hesitation locations in the Risk Perception Tests (Site 1 – Fixed / Site 2 – Risky). Here, Site 1 was associated with a fixed 150s hold time and Site 2 with a 50% chance of either a 1s or 300s hold time. During training Trips 1–20, foragers (n = 8) were collected after they exited the testing grid (grey line, Collection Point) at both Site 1 and 2 (8m and 6m from the nest entrance respectively). During the Test Trip, foragers were tested with no corresponding homeward vector (Zero Vector) and allowed to pass through both sites and travel to the nest entrance.

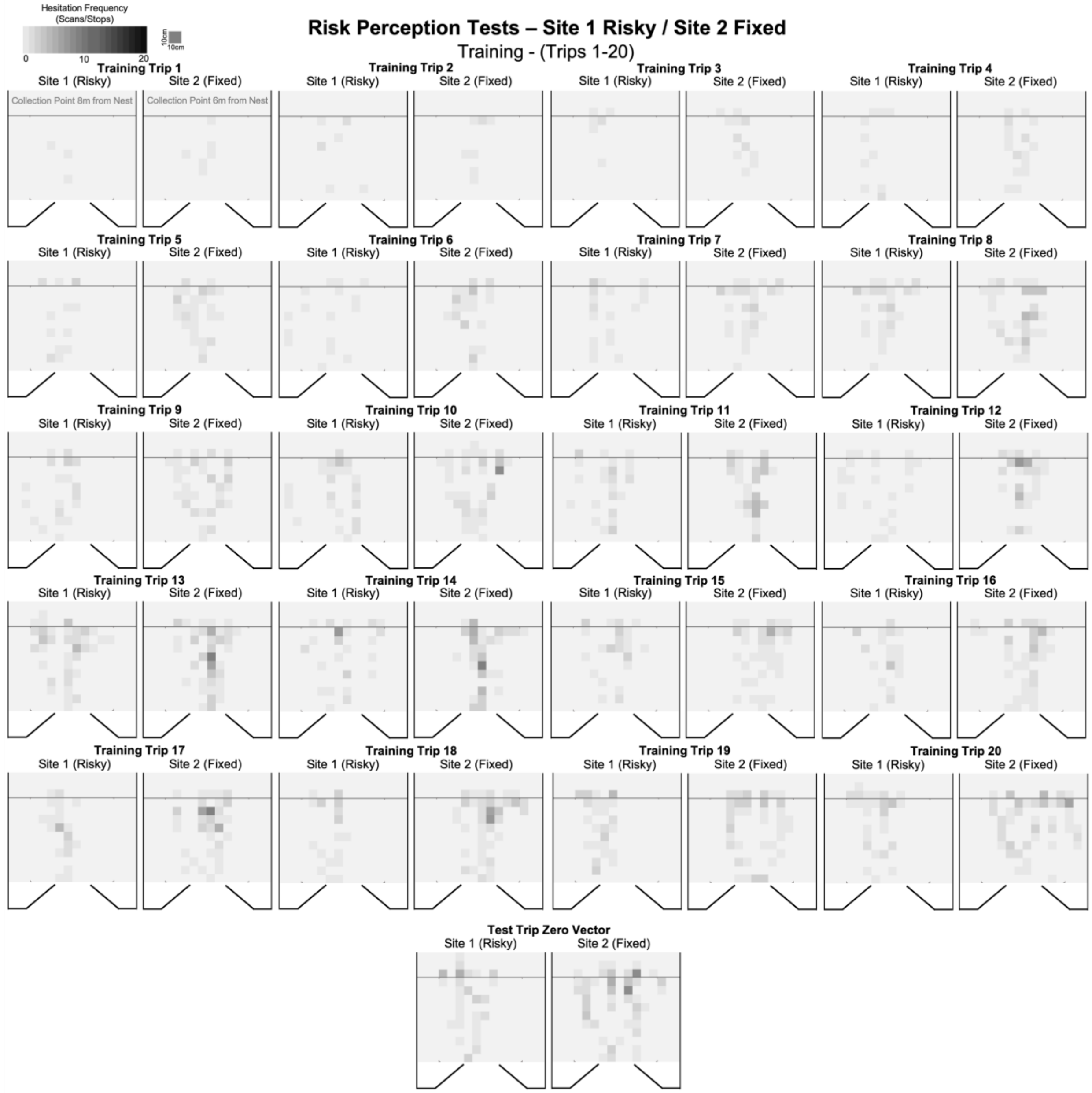


**SFig. 5.** Heat maps of forager hesitation locations in the Risk Perception Tests (Site 1 – Risky/ Site 2 – Fixed). Here, Site 1 was associated with a 50% chance of either a 1s or 300s hold time and Site 2 was associated with a fixed 150s hold time. During training Trips 1–20, foragers (n = 7) were collected after they exited the testing grid (grey line, Collection Point) at both Site 1 and 2 (8m and 6m from the nest entrance respectively). During the Test Trip, foragers were tested with no corresponding homeward vector (Zero Vector) and allowed to pass through both sites and travel to the nest entrance.
